# Structural Coordinates: A novel approach to predict protein backbone conformation

**DOI:** 10.1101/2020.09.15.297747

**Authors:** Vladislava Milchevskaya, Alexei M. Nikitin, Sergey A. Lukshin, Ivan V. Filatov, Yuri V. Kravatsky, Vladimir G. Tumanyan, Natalia G. Esipova, Yury V. Milchevskiy

## Abstract

**Motivation:** Local protein structure is usually described via classifying each peptide to a unique element from a set of pre-defined structures. These so-called structural alphabets may differ in the number of structures or the length of peptides. Most methods that predict the local structure of a protein from its sequence rely on this kind of classification. However, since all peptides assigned to the same class are indistinguishable, such an approach may not be sufficient to model protein folding with high accuracy.

**Results:** We developed a method that predicts the structural representation of a peptide from its sequence. For 5-mer peptides, we achieved the Q16 classification accuracy of 67.9%, which is higher than what is currently reported in the literature. Importantly, our prediction method does not utilize information about protein homologues but only physicochemical properties of the amino acids and the statistics of the structures, but relies on a comprehensive feature-generation procedure based only on the protein sequence and the statistics of resolved structures. We also show that the 3D coordinates of a peptide can be uniquely recovered from its structural coordinates, and show the required conditions for that under various geometric constraints.

**Availability:** The online implementation of the method is provided freely at http://pbpred.eimb.ru

**Supplementary information:** Supplementary data are available online at http://pbpred.eimb.ru/S/index.html

## Introduction

Due to fast development of machine learning and statistical inference methods, recent years brought significant progress in protein local structure classification and prediction. [1, 2] The predicted structure of local protein fragments serves as a critical component in most global protein folding models, along with homology information and physicochemical properties of amino acids. [3]

The majority of local structure prediction methods rely on the classification of these fragments into certain structural classes, also referred to as structural alphabet or protein blocks.[4]. One of the widely used methods, DSSP, exploits hydrogen bond information to cluster protein 5-mers into eight structural classes that include *α*-helices, *β*-sheets, 3/10 helices et.c.[5] Frischman et al. extend this by incorporating the quantitative information of the dihedral angle frequency into their prediction algorithm STRIDE.[6]

Other methods calculate a distance matrix between *C_α_* atoms of the backbone a fragment and use the cumulative distance between the corresponding matrix elements to measure dissimilarity between two protein fragments. Based on this, the authors proposed a structural alphabet containing 27 clusters of 7-mer fragments.[7] Hahn et al. first cluster fragments based on their sequence similarity, and then examine structural variation within clusters.[8] The ones with high within-cluster variation are discarded, while others are kept to represent sequence neighbourhoods.

One of the most used structural alphabets at the moment has been developed by de Brevern et al. [9] There, the authors clustered all five-mer peptides available in the PDB at that time into 16 groups based on the RMSDA (root mean square deviation of angle) distance between the peptides (1). The authors refer to this alphabet as protein blocks (PB).

Joseph et al. review various applications of the Protein Blocks (PB): Local protein structure prediction, structural alignment, identification of functional structural motifs in proteins et.c.[2]

However, these and most other methods for local structure prediction available in the literature are classification-based. Namely, a set of structural clusters is given, and each protein fragment is assigned a unique cluster label. Consequently, if two protein fragments were assigned the same cluster label, they can not be distinguished from one another in three-dimensional coordinates (3D). However, there is significant variation in structure within clusters, which is why classification-based approaches may not be sufficient to model protein folding at high resolution.

Here we propose a new approach for using alphabets of basic structures, and represent each peptide in terms of distances to these structures (structural coordinates). We show that this approach yields better classification and prediction accuracy than the current gold-standard. Additionally, we show that given enough basic structures, the 3D coordinates of a peptide can be uniquely recovered from its structural coordinates.

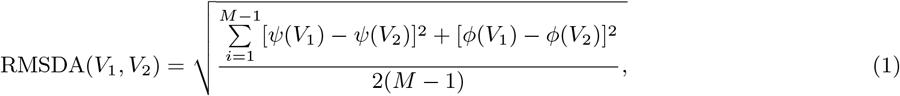

*where M is the number of residues in a fragment (in the original article M* = 5), *V*_1_ *and V*_2_ *denote the fragments, and* {*ψ_i_*(*V*_1_), *ϕ_i_*(*V*_1_)}, *i* = 1,…, 2(*M* − 1) *denote the series of dihedral angles for fragment V*_1_.

## Results

### Description of protein clusters in terms of RMSD

For local structure classification and prediction, one of the most used sets of protein structures is a set of the 16 protein blocks (PB) presented by De Brevern et al. [9]. These PBs serve as cluster centres: A given protein structure is classified according to its nearest protein block and assigned the corresponding cluster label. In the original article, the authors use the RMSDA (root mean square deviation of angular values) quoted in (1) to define the distance between two protein structures. Here we use the same PBs as cluster centres but compose clusters out of the nearest in terms of RMSD protein structures from our training sample. Table 1 shows locations frequencies of the secondary structure, with respect to our training sample. As expected, the coarse characterization of the clusters agrees with the one presented by de Brevern et al. Namely, for the clusters that describe the major regular structures, such as *α–*helices and *β–*strands, the relative size of the clusters differs only slightly between RMSD- and RMSDA-based compositions. However, for the clusters marked as “mainly coil”, the differences are more pronounced.

**Table 1:**
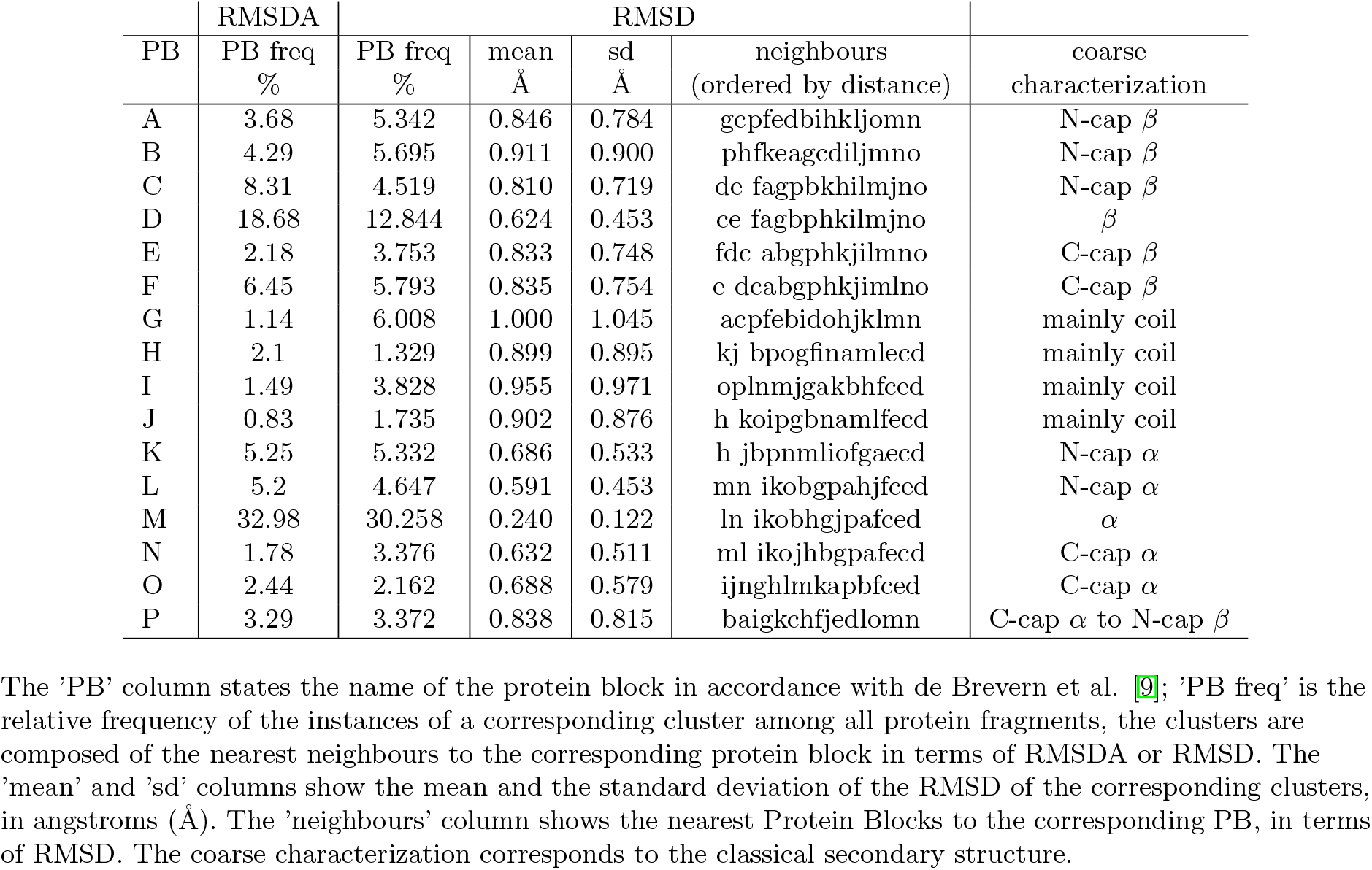
Description of protein clusters in terms of RMSD

On the contrary, the composition of the RMSD- and RMSDA-based clusters appears to have noticeable differences: In total, only 69.6% of the 5-residue fragments from the training sample have identical labels in both cluster sets. Table S1 and Fig. S1 (online) show how cluster composition varies for the two distance measures in more details. Thus, cluster *m* which corresponds to *α*–helical structures is almost identical in both cases (97.4%). The same holds for the clusters *l* and *n* that are structurally similar to *m* (see Tab. S2 (online)). Similarly, RMSDA-based clusters *c, d* and *e*, that cover *β*-sheet-like structures, agree well with the corresponding RMSD-based ones: *C, D* and *E*. Note here that *PB_c_, PB_d_* and *PB_e_* structures are very similar to one another in terms of RMSD, which is why the high percentage of confusion between the *d* and *E* or any other discrepancy within this group is expected.

We observed that in general, the more regular is the structure described by a certain cluster, the more similar is the composition of that cluster for both distance measures. The largest discrepancy in RMSD- and RMSDA-based cluster composition appears for those clusters that do not cover regular or partially regular structures: Namely, those marked “mainly coil” in Tab. 1. For instance, cluster *G* only has 8.7% of its structures classified as *g,* while the remaining structures get various labels of a number of other clusters (*c,d,f,l,m*). Notably, cluster G has the largest mean and standard deviation of RMSD from its members to the cluster centre *PB_g_*: 1.00 and 1.04 angstroms, while for a compact *α*-helical cluster *M* it is 0.24 and 0.12 angstrom correspondingly. This notion may present a significant challenge for interpretation of the structures that belong to such a non-compact (high variance and large mean RMSD to the central structure) cluster, as well as hamper the prediction accuracy.

### RMSD recovers ambiguity of RMSDA

Dihedral angles of a protein fragment represent one of the most important characteristic of the structure, and researchers often rely on this characteristics to classify structures. For instance, the standard alpha-helical fragment of five residues would have the following dihedral angles: *ϕ_i_* = −57, *ψ_i_* = −47, *i* = 1,…,5. However, two protein fragments may have the same RMSDA to the standard helical structure but their actual alignment in 3D may differ significantly. Fig. 1 shows four protein fragments of 5 residues (in green) aligned to the standard helical fragment (red), RMSDA of the green and the red structures are the same for all four cases. Yet, the dissimilarity between the standard helix and the first fragment is very small, whereas the second and the third fragments are noticeably different from the standard alpha-helix. The fourth fragment represents a rare case of the peptide bond angle ω ≠ 180: in terms or RMSDA it is identical to the alpha-helix, while the two structures are considerably different. These examples illustrate that structural similarity between protein fragments is not captured by the RMSDA measure entirely. Another situation in which RMSDA may under- or overestimate the similarity between two three-dimensional fragment may be, for instance, if all their dihedral angles are identical except for one angle in the middle (for instance, *ψ*_3_ in a 5-residue fragment): These structures will not align close, even though their RMSDA may be small. Based on these considerations, we chose to use RMSD as distance measure between protein fragments over RMSDA proposed in the original article. [9]

**Figure 1:**
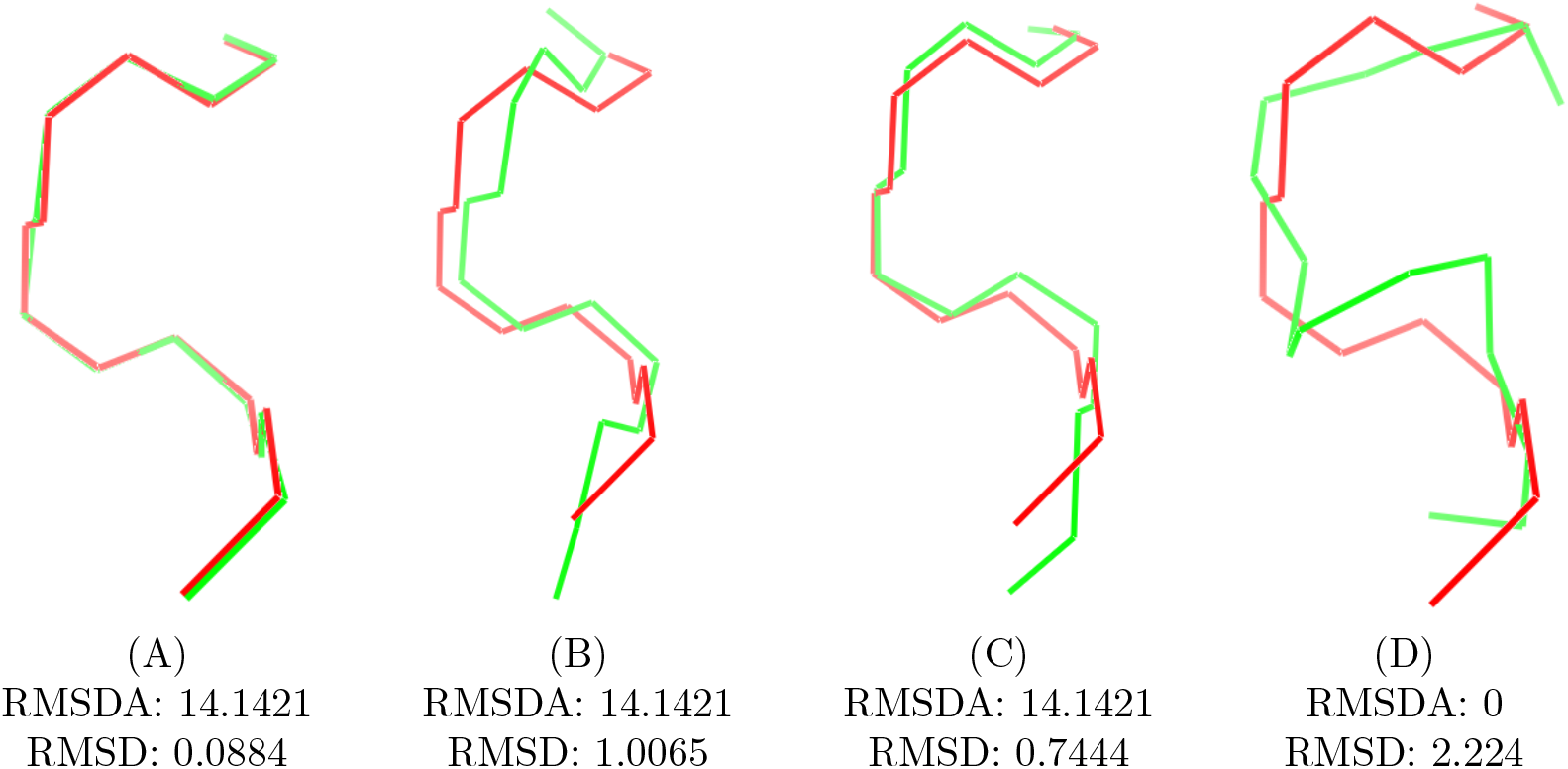
Structural alignment of 5-residue protein fragments (green) against the standard alpha-helical structure (red) with dihedral angles *ϕ_i_* = −57, *ψ_i_* = −47, *i* = 1,…, 5. Dihedral angles of the fragment (A) are *ϕ_i_* = −61.142, *ψ_i_* = −42.857, *i* = 1,…, 5, its RMSDA to the standard helix is 14.1421 and its RMSD to the standard helix is 0.0884; Dihedral angles of the fragment (B) are *ϕ_i_* = −32.857, *ψ_i_* = −42.857, *i* =1,…, 5, its RMSDA to the standard helix is 14.1421 and its RMSD to the standard helix is 1.0065; Dihedral angles of the fragment (C) are all *ϕ_i_* = −57, *ψ_i_* = −47, except for *ϕ_3_* = −17, its RMSDA to the standard helix is again 14.1421 and its RMSD to the standard helix is 0.7444. Dihedral angles of the fragment (D) are all *ϕ_i_* = −57, *ψ_i_* = −47, except *ω*_3_ =0 instead of the usual 180.

### Reconstruction of the backbone conformation

There are several advantages of measuring similarity between protein structures based on RMSD. First of all, RMSD allows us to recover dihedral angles used in RMSDA utilized in previous works. [9] Furthermore, given enough basic structures, one can also reconstruct 3D coordinates of a protein fragment from its RMSD-distances to basic structures, which can not be achieved by using RMSDA. We illustrate this in the following examples.

#### Recovering angles

First of all, knowing RMSD between a protein structure and the 16 basic protein blocks allows us to recover dihedral angles of that structures. To illustrate this, we fixed a 5-residue fragment *x* with the dihedral angles

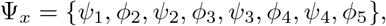

there actual values are shown in Table S4 (online). Let its distances to basic structures *PB*_1_ *,PB*_2_,…, *PB*_16_ be denoted as 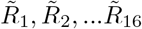 correspondingly. For this example, *R_i_* can be seen as a function of dihedral angles, i.e *R_i_* = *R_i_* (Ψ), *i* = 1,…,16.

To reconstruct the dihedral angles Ψ_*x*_ of the target structure *x*, we minimised the squared loss as a function of angles:

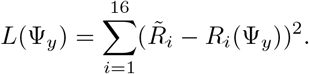

For that, we used two optimization methods: The method of conjugate gradients and the Fletcher-Powell algorithm, the starting point for both algorithms was set to Ψ_0_ = (0, 0, 0, 0, 0, 0, 0, 0). Table 2 shows that both methods reached the true values of dihedral angles with high accuracy.

**Table 2:** Predicted and actual angles

|  | True angles | Conj. Grad. | Fletcher-Powell |
| --- | --- | --- | --- |
| $\psi_1$ | 108.239 | 108.048 | 108.240 |
| $\phi_2$ | -90.119 | -90.011 | -90.120 |
| $\psi_2$ | 119.539 | 119.493 | 119.540 |
| $\phi_3$ | -92.209 | -91.988 | -92.210 |
| $\psi_3$ | -18.059 | -18.458 | -18.060 |
| $\phi_4$ | -128.929 | -128.636 | -128.930 |
| $\psi_4$ | 147.039 | 147.369 | 147.041 |
| $\phi_5$ | -99.899 | -100.416 | -99.901 |

Predicted and actual dihedral angles of the protein fragment $x$ . The 'True angles' are the actual dihedral angles of the protein fragment $x$ ; the 'Conjugate Gradients' and 'Fletcher-Powell' show the angle values reconstructed from the distances to the 16 protein blocks using the conjugate gradients and Fletcher-Powell algorithms correspondingly.

### Number of PB needed for reconstruction

We have seen that with 16 basic structures (protein blocks) it is possible to reconstruct the backbone coordinates of a 5-residue protein fragment, given the bond lengths and most of the bond angles are fixed. Here we calculate the minimal number of basic structures required in the general case. Let us consider a N-residue fragment, each amino acid corresponds to three backbone atoms (*N,C_α_,C*), and each atom has three coordinates in 3D. Thus, to define the position of the backbone, one needs 9*N* degrees of freedom. Further, if the backbone is shifted along *X,Y* or *Z* axis, the structure is unchanged – thus, one can omit the 3 translational degrees of freedom. Similarly, the three rotational degreed of freedom do not represent the protein structure, and can be omitted as well. In most proteins, the change in bond lengths can be neglected – therefore, one may subtract 3*N* − 1 degrees of freedom. Fixing bond angles would subtract additional 3*N* − 2 degreed, and setting the *ω* angle (peptide bond) to 180 would release another *N* − 1 degrees of freedom. Tab. 3 sums up these considerations.

**Table 3:** Degrees of freedom

| Geometric constraints | deg. of freedom |
| --- | --- |
| Positions of all $3N$ atoms | $9N$ |
| Rotation and Translation invariance | $9N - 6$ |
| Bond lengths are fixed | $6N - 5$ |
| Bond angles are fixed | $3N - 3$ |
| Peptide bond angles $\omega$ are fixed | $2N - 2$ |

The minimal number of degrees of freedom required to describe an $N$ -residue protein backbone, given a set of geometric constraints.

Therefore, for a 5-residue fragment with all bond length and bond angles considered fixed, one would need at least 12 (fairly unrelated) basic structures to reconstruct the conformation of the backbone. With the same restrictions for a 9-residue fragment, at least 24 basic structures would be required.

### Benchmarking Q16

Here we return to the classification problem and benchmark our method against the two leading tools for protein local structure prediction: LOCUSTRA and PB-kPred. [10, 9] In order to achieve comparable figures, we use the same protein fragment size (5 residues) and the same set of proteins as in PB-kPred (PDB30) for our model training and testing.

Table S3 (online) shows that for clusters *B, D, F, G, J, K, M, O* and P out method has higher prediction accuracy than the two other methods, whereas for clusters *A, C, H* and *N* PB-kPred shows better performance. Overall, the accuracy of our Q16 classification is 67.9% which is 6% higher than the one reported for PB-kPred[11]. Importantly, unlike the two other prediction methods, our model does not rely on the information about homologous proteins. Therefore, the distribution of prediction accuracies for our method is unimodal Gaussian, but not bimodal as for PB-kPred, where the high prediction accuracy corresponds to those proteins that have homologs outside the training sample (see Fig. S3 online.).

## Materials and methods

### Structure Database generation

#### Datasets for training and testing

The PDB contains a high number of nearly identical molecules, which may add undesired confounding factors at the model training step. To alleviate this, we used a representative set of sequence-unique structures with high resolution. Furthermore, to ensure that our results are comparable to those from the PB-kPRED prediction method, we opted for the PBD30 dataset comprised by Vriend et al. [11] The PBD30 dataset contains 15,544 protein chains restricted to 30% sequence identity according to BLASTclust characterisation. [12] Among the sequences that shared more than 30% identity, the researches chose one representative protein chain that corresponds to the best available structure. Out of the 15,544 structures, over 14000 were crystallographic, approximately 1100 were NMR-resolved, and 209 were solved by electron microscopy. The full list of the PDB identifies can be found at http://pbpred.eimb.ru/S/LS.txt.

The training and testing sets were obtained by partitioning the full set of 15,544 protein fragments into five equally sized subsets. Further, four of these subsets are used for training, and the remaining subset is used for testing.

#### Dissimilarity measure

To classify each of the pentapeptides from the training set, we use the 16 protein blocks (PB) developed by de Brevern et al., also referred to as Q16. [9] Since PBs have the same length of five residues, one can calculate RMSD (root mean square deviation) between a pentateptide and a protein block *PB_i_, i* = 1,…, 16.

The smaller the RMSD is, the closer the two protein fragments are in terms of structure. Further, we define a similarity measure:

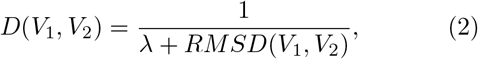

where λ is a fixed constant such that λ ≥ 0. Here large values of *D* correspond to more structurally similar fragments. To assign a five-mer protein fragment to one of the Q16 classes, we calculate its similarity value to each of the 16 protein blocks *PB_i_, i* = 1,…, 16.

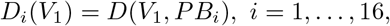

and choose the class with the highest similarity value.

### Structural alignment

Computation of the RMSD between two protein fragments requires optimisation over all possible alignments of these fragments against each other. Kabsch et al. introduced an optimisation procedure and showed that it reached the minimum of the RMSD function.[13, 14] We adapted a version of this procedure implemented by Gans and Shalloway in their tool *Qmol,* and integrated it into our pipeline. [15]

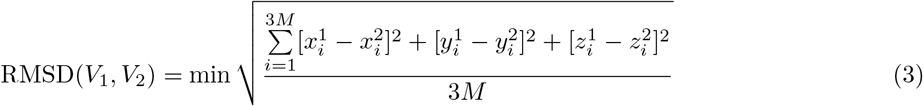

*where M is the number of residues in a fragment (in the original article M* = 5), *V*_1_ *and V*_2_ *denote the fragments, and* 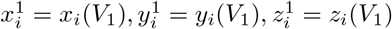, *i* = 1,…, 3*M denote Cartesian coordinates of the backbone atoms of the M–residue fragment V*_1_, *and the minimum is taken over all spatial co-localization of the two fragments V*_1_ *and V*_2_.

### Feature generation

Generation of a comprehensive pool of features is the main and most laborious step of the prediction algorithm.

The sequence of a protein fragment can be characterised by the properties of amino acids forming this fragment as well as its flanking regions. At the model selection step, we tried flanking regions of various length, including the complete protein sequence. Two main types of information were used to generate a predictor pool: Physico-chemical properties of amino acids and RMSD distribution of fragments with the identical sequence to each of the protein blocks *PB_i_, i* = 1,…, 16.

#### RMSD-based predictors

reflect the distribution of distances from a given sequence fragment to the 16 protein blocks. These distributions are derived from the identical sequence instances in the training set. Unsurprisingly, we encountered the following problem: For sequence fragments of five amino acids, there are 20^5^ = 3200000 possible letter combinations. Thus, even for five-residue peptides, certain sequence combinations are massively underrepresented in the training set, which makes it impossible to acquire reliable statistics on their structure.

To alleviate this, we developed so-called reduced amino acid alphabets, where certain amino acids are seen as identical. In this definition, identical sequence fragments can be described as regular expressions. For instance, all aliphatic amino acids ([GAVLI]) or sulfur-containing amino acids ([CM]), or all aromatic amino acids ([YWF]) may be assigned a single identity class. These rules may differ based on the position of the amino acid inside a protein fragment, because for instance, the central amino acid may have a stronger influence on the predicted structure. All reduced amino acid alphabets are available in the Supplementary via http://pbpred.eimb.ru/S/reduced_alphabets.zip

After sequence identity classes are defined, we calculate the minimal sufficient statistics for the distance distribution of each identity class. Namely,

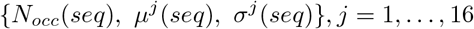

where *N_occ_*(*seq*) is how many times instances of the sequence seq occur in the training sample, *μ^j^*(*seq*) and σ^j^ (*seq*) are the mean and variance estimates of the RMSD between the fragments from the current identity class and the *j*–th protein block. Further, we applied various non-linear transformations to generate a pool of predictors. Fig.2 schematically represents the predictor generation procedure. One example of an RMSD-based predictor a t-statistics-like function, which captures differences between an identity of a certain sequence class and the whole sample. This and a more detailed description of the generation procedure can be found in the Supplementary (online).

**Figure 2:**
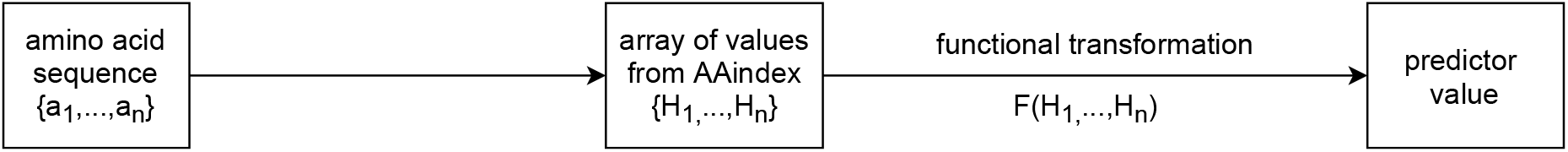
Generation of predictors based on RMSD statistics between a protein fragment and the 16 basic protein blocks, schematic representation.

#### Physico-chemical properties

of amino acids were acquired from the AAindex database [16]. Among the 550 different properties present in the database, we used various scales of hydrophobicity, flexibilities of amino acid residues, solvation free energy et.c. We will illustrate predictor generation with an example: Assume, there is a sequence fragment of n residues {a_1_,…,a_n_}. Each amino acid {*a_i_, i* = 1,…*,n*} has a certain value of hydrophobicity {*H_i_,i* = 1,…*,n*}. Further, a functional transformation was applied to the array of property values {*H*_1_,…, *H_n_*}, i.e. F(H_1_,…, *H_n_*). In the current example, the functional transformation reflects a periodic change of hydrophobicity with period *T* in a given fragment of n residues:

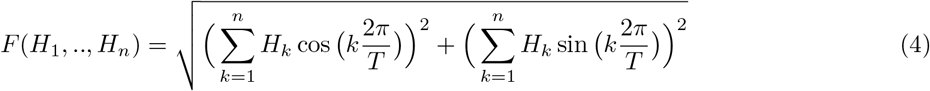

For other physico-chemical properties, functional transformations may be different. Fig.3 illustrates the general procedure for generating this kind of predictors. Note that at this step, no alphabet reduction was used.

**Figure 3:**
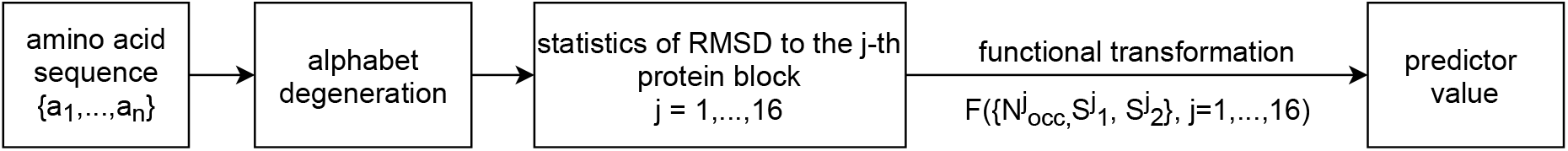
Generation of predictors based on physico-chemical properties of amino acids comprising a protein fragment, schematic representation.

### Model fitting

Further, we train a linear regression model to estimate the similarity *D_i_*(*V*) (2) between a protein fragment *V* and each of the protein blocks *PB_i_*. Assume 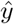 is the estimate of *D_i_*(*V*) for some *i* = 1,…, 16, and {*x*_1_,…, *x_p_*} are all the predictor values generated above. Then,

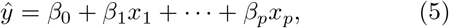

where *β*_0_,…, *β*_p_ are the regression coefficients. In order to fit the regression coefficients, we adapted the method of stepwise linear regression, with bidirectional elimination of predictors. Model fitting procedure is described in details in the Supplementary (online).

## Discussion

The rapidly increasing amount of information about protein structures, coupled with advances in machine learning, opens many possibilities to tackle the problem of protein structure prediction. One of the common limiting factors in local structure prediction is the availability of homologues to the target protein in the PDB. Another challenge is recovering the coordinates of the protein backbone given its sequence.

In this work, we introduced a novel approach to predicting local structure that does not rely on information about protein homologues. It is particularly useful for predicting the local structure of those proteins that do not have any homologues in the PBD. Mainly, we decomposed a complex multidimensional problem of protein stricture prediction into several one-dimensional regression tasks. Each regression problem aims to quantify the relationship between a protein sequence, the statistics of its structural conformations and the physicochemical properties of its amino acids.

Using an appropriate feature space is one of the keys to achieve high prediction accuracy for almost every leaning algorithm. We believe, the main contribution of our method is the generation of a comprehensive pool of features adapted for various challenging scenarios.

In particular, rare sequence fragments posed a significant challenge for training our model. To tackle this, we developed a so-called reduced alphabet: Sequence fragments were mapped to specific regular expressions which took physicochemical properties of the amino acids into account to form new sequence identity classes. This way, we achieved more abundant identity classes with presumably similar conformation properties.

Another set of the features with high predictive power relies on the physical properties of the structural fragments. For instance, predictor (4) reflects the periodic change of hydrophobicity. When the periodicity value *T* = 3.6, it corresponds to a feature with the highest predictive power for alpha-helical conformation (Protein Block ‘m’). Thus, when the value of this feature is high, hydrophobic and hydrophilic residues are likely to face opposite direction, which is energetically favourable for a helical fragment on the surface of a protein.

To benchmark our method against the current gold standard, we used the same set of proteins for training and testing as in PB-kPred, and the same set of 16 basic structures (protein blocks). We achieved the Q16 classification accuracy of 67.9% independent of the availability of homologues, which is higher than the 40.8% to 66.3% depending on the availability of homologues by PB-kPred.[9]

One major difference between our method and PB-kPred is that RMSD but not RMSDA was used to measure the distance between two protein fragments. Although the calculation of RMSD is computationally more demanding, we have shown that for local structure prediction, RMSD measure has considerable advantages. Structures that are close in terms of RMSDA may not align well in 3D, while for RMSD it always holds. Further, we have split all 5-residue protein structures into 16 clusters based on the nearest protein block in terms of RMSD. The clusters that corresponded to regular structures (alpha-helix, beta-sheet) had a high overlap with those based on RMSDA. In contrast, clusters with variable structural composition differed between the two distance measures.

Further, as a proof of concept, we reconstructed the structures of several protein fragments solely from the (actual) RMSD values between a fragment and the 16 protein blocks. Using the notion of degrees of freedom, we established the lowest number of basic structures required to reconstruct the conformation of a protein fragment of a given length, with various geometric constraints. In practice, the reconstruction may benefit from a larger number of basic structures since distances to protein blocks are predicted and contain a certain amount of noise. In future works, we aim to determine the optimal set of basic structures and their effective number via clustering based on RMSD. This way, the protein blocks will be most variable in terms of the same metric as the one used in the regression model.

The small fraction of membrane and NMR-resolved proteins structures in the utilised dataset may not be explained well by our prediction model. Garbuzynskiy et al. show that NMR-resolved structures have systematic differences form the protein structures resolved by X-ray. [17] This batch effect needs to be accounted for if the structures from both methods are to be used together. Additionally, the connection between a protein structure and its physicochemical properties varies drastically between the membrane and non-membrane proteins. Therefore, separating these tasks from one another and pre-filtering the training set may improve the prediction accuracy of the model.

The proposed approach can be further extended to several scenarios: Fast PDB mining for similar structures using predicted distances to the basic protein blocks. In molecular dynamics, researchers use predicted structures of small protein fragments as an initial approximation and constraints for the model. To this end, we believe that protein blocks of different length may be needed and utilised together.

## Acknowledgements

We thank Alexej Viktorovich for the valuable input during the manuscript preparation.

