## Supplementary for "Structural Coordinates: A novel approach to predict protein backbone conformation"

### S1 Predictor generation and examples

One of the two main sources for feature generation is the statistics of occurrence of a certain protein sequence in the sequences of other proteins with resolved structures. Here we combine it with the RMSD distance to the sixteen protein blocks.

**A predictor based on the t-statistics.** For instance, let us consider one of the sixteen protein blocks $PB_j, j \in \{1, 2, 3, \ldots, 16\}$, and a 5-residue sequence $seq$. Also, let $N_{occ}(seq)$ be the number of times sequence $seq$ occurs among the sequences with known structures (the training sample), $\bar{\mu}_j = \bar{\mu}_j(seq)$ be the mean distance between the structures with that sequence and the $PB_j$. Further, let $\bar{\mu}_j$ be the average distance between $PB_j$ and all 5-residue fragments in the training sample, and $s_j^2$ be its sampling variance and $N$ to be the size of the training sample. Then, one example of the predictors is:

$$t_j(seq) = \frac{\mu_j - \bar{\mu}_j(seq)}{s_j(seq)},$$

where

$$s(seq) = \frac{\sigma_j^2(seq)}{N_{occ}(seq)} + \frac{s_j^2}{N}.$$

Note that when $N \gg N_{occ}(seq) > 1$, $s \cong \frac{\sigma_j^2(seq)}{N_{occ}(seq)}$, the following holds:

$$t_j(seq) \cong \frac{\mu_j - \bar{\mu}_j(seq)}{\sigma_j(seq)} \sqrt{N_{occ}(seq)}.$$

Thus, the number of occurrences of as certain sequence in the sample, $N_{occ}(seq)$ is crucial for correct estimation of $t_j(seq)$. Namely, small values of $N_{occ}(seq)$ may yield unreliable estimates of $\sigma_j(seq)$ as well as $t_j(seq)$. To alleviate this, we tried various reduced alphabets.

A reduced alphabet is defined by the identity classes of the amino acids. For instance, in the following reduced alphabet example (tab.S4) for a seven-residue protein sequence all amino acids are different in positions -3, -2, 1 and 2. In positions -1, 0 and 1 amino acids $\{A, L, M, C\}$ are indistinguishable and make one identity class, amino acids $\{G, P\}$ make another identity class, $\{V, I, F, Y, W, K, R, H, D, E, N, Q, S, T\}$ make the third identity class, $\{O\}$ makes the fourth and $\{X\}$ makes the fifth:

Table S4: **An example of reduced alphabets for a 7-residue sequence fragment**

| | |
|---|---|
| -3 | A V L I P M C F Y W K R H D E N Q S T G O X |
| -2 | A V L I P M C F Y W K R H D E N Q S T G O X |
| -1 | ALMC VIFYWKRHDENQST GP O X |
| 0 | ALMC VIFYWKRHDENQST GP O X |
| 1 | ALMC VIFYWKRHDENQST GP O X |
| 2 | A V L I P M C F Y W K R H D E N Q S T G O X |
| 3 | A V L I P M C F Y W K R H D E N Q S T G O X |

### S2 Model fitting

Here we fit a linear regression model to estimate the similarity $D_i(V)$ (2) between a protein fragment $V$ and each of the protein blocks $PB_i$. Thus, each protein fragment $V$ (we mostly use 5-residue long fragments) corresponds to 16 values $\{D_i(V)\}_{i=1,\dots,16}$. The fit is performed independently for each of the 16 components $D_i(V)$.

Assume $\hat{y}$ is the estimate of $D_i(V)$ for some fixed $i = 1, \dots, 16$, and $\{x_1, \dots, x_p\}$ are all the predictor values generated as described before. Then the model takes the following form:

$$\hat{y} = \beta_0 + \beta_1 x_1 + \cdots + \beta_p x_p, \tag{6}$$

where $\beta_0, \dots, \beta_p$ are the regression coefficients. Commonly, the regression coefficients are estimated to minimise the squared error loss function. However, here $p$ is large and many predictors will not be meaningful. Additionally, a number of predictors among $\{x_1, \dots, x_p\}$ will be correlated due to the generation procedure. Hence, we first use stepwise forward-backward selection procedure to screen for the "good" predictors that show fairly high predictive capability (in terms of F-statistics), and then fit the regression model for this decreased set of predictors via least squares. The threshold for the F-statistics is determined via leave-one-out cross-validation on the training subset of the data.

### S3 Tables and Figures

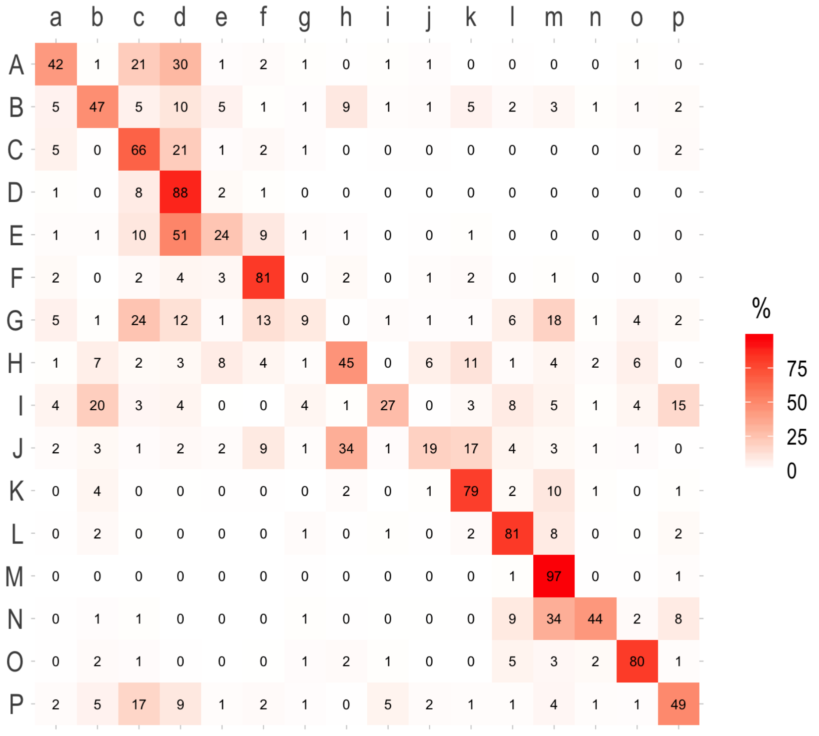


Figure S4: Cluster assignment discrepancy. Percentage of structural fragments in the learning sample shared between RMSD- and RMSDA- based clustering of those fragments.

Table S5: **Cluster assignment discrepancy: RMSD vs. RMSDA**

| | a | b | c | d | e | f | g | h | i | j | k | l | m | n | o | p |
|---|---|---|---|---|---|---|---|---|---|---|---|---|---|---|---|---|
| A | 42.4 | 0.7 | 20.7 | 29.7 | 0.8 | 1.6 | 0.7 | 0.3 | 0.7 | 0.9 | 0.1 | 0.5 | 0.1 | 0.1 | 0.6 | 0.3 |
| B | 4.9 | 47.2 | 5.4 | 10.5 | 4.9 | 0.7 | 0.9 | 9.1 | 1.2 | 1.4 | 4.9 | 1.9 | 3.3 | 0.7 | 1.4 | 1.7 |
| C | 5.3 | 0.5 | 65.7 | 20.7 | 1.3 | 2.4 | 0.9 | 0.1 | 0.3 | 0.3 | 0.1 | 0.1 | 0.1 | 0.1 | 0.3 | 1.8 |
| D | 0.6 | 0.4 | 8 | 87.6 | 2.3 | 0.7 | 0 | 0 | 0.1 | 0.1 | 0.1 | 0 | 0 | 0 | 0 | 0.1 |
| E | 1 | 1.2 | 9.8 | 51 | 23.8 | 9.1 | 1.4 | 0.9 | 0.1 | 0.4 | 0.7 | 0 | 0 | 0.1 | 0.1 | 0.3 |
| F | 2.3 | 0.1 | 1.9 | 3.6 | 3.3 | 81.1 | 0.5 | 2.3 | 0 | 1.1 | 1.9 | 0.5 | 0.7 | 0 | 0.3 | 0.4 |
| G | 5.4 | 0.7 | 24.4 | 12.3 | 1.4 | 12.6 | 8.8 | 0.3 | 1.4 | 0.9 | 0.9 | 6.2 | 18.2 | 0.6 | 3.9 | 2.1 |
| H | 0.6 | 6.6 | 2.1 | 2.8 | 7.5 | 3.7 | 1.1 | 44.9 | 0.1 | 6.2 | 11.2 | 1.4 | 3.8 | 1.5 | 6.1 | 0.4 |
| I | 4.4 | 20 | 3.4 | 4.1 | 0.2 | 0.1 | 3.7 | 0.7 | 27.4 | 0.5 | 2.6 | 8.4 | 4.7 | 0.6 | 3.8 | 15.3 |
| J | 1.6 | 2.8 | 1.3 | 2.4 | 1.5 | 8.7 | 1.1 | 33.6 | 1.3 | 18.7 | 17.1 | 4.3 | 2.8 | 1.1 | 1.3 | 0.4 |
| K | 0.1 | 3.6 | 0.3 | 0.2 | 0.4 | 0.4 | 0.1 | 1.6 | 0.1 | 0.6 | 79.4 | 2.3 | 9.6 | 0.6 | 0.1 | 0.7 |
| L | 0.5 | 2.4 | 0.2 | 0 | 0 | 0 | 1.4 | 0.5 | 0.6 | 0.5 | 1.9 | 81.1 | 8.1 | 0.5 | 0.2 | 2.3 |
| M | 0 | 0 | 0 | 0 | 0 | 0 | 0.3 | 0 | 0 | 0 | 0 | 0.6 | 97.4 | 0.3 | 0.3 | 1 |
| N | 0.4 | 0.8 | 0.6 | 0.1 | 0 | 0.1 | 0.6 | 0.4 | 0.2 | 0.2 | 0.5 | 8.8 | 33.6 | 43.5 | 2 | 8.1 |
| O | 0.5 | 1.9 | 1.3 | 0.5 | 0.1 | 0.2 | 1.3 | 2 | 0.6 | 0.1 | 0.1 | 4.8 | 2.9 | 2.2 | 80.4 | 1 |
| P | 1.5 | 4.9 | 16.9 | 9.2 | 1 | 1.9 | 0.8 | 0.3 | 4.7 | 1.5 | 1.2 | 1.1 | 3.5 | 1.2 | 0.9 | 49.3 |

The rows correspond to the RMSD-based clusters, and the columns correspond to the RMSDA-based clusters. As before, the clusters are formed out of the fragments nearest to the corresponding protein block, in terms of RMSD (capital letters) or RMSDA (small letters). The values in the represent the percentage of fragments shared by a pair of clusters: For instance, clusters 'm' and 'M' share 97.4% of fragments.

Table S6: **Distances between cluster centres**

| | a | b | c | d | e | f | g | h | i | j | k | l | m | n | o | p |
|---|---|---|---|---|---|---|---|---|---|---|---|---|---|---|---|---|
| a | 0 | 1.58 | 1.28 | 1.56 | 1.51 | 1.44 | 1.03 | 2.16 | 1.85 | 2.23 | 2.19 | 2.23 | 2.37 | 2.5 | 2.25 | 1.41 |
| b | 1.58 | 0 | 1.77 | 1.88 | 1.57 | 1.54 | 1.7 | 1.53 | 2.02 | 2.05 | 1.57 | 2.02 | 2.07 | 2.12 | 2.52 | 1.37 |
| c | 1.28 | 1.77 | 0 | 0.65 | 0.96 | 1.15 | 1.52 | 2.65 | 2.71 | 2.93 | 2.63 | 2.79 | 2.81 | 3.03 | 3.13 | 1.73 |
| d | 1.56 | 1.88 | 0.65 | 0 | 0.81 | 1.14 | 1.87 | 2.79 | 3.02 | 3.17 | 2.84 | 3.07 | 3.12 | 3.34 | 3.49 | 2.02 |
| e | 1.51 | 1.57 | 0.96 | 0.81 | 0 | 0.61 | 1.63 | 2.37 | 2.8 | 2.8 | 2.5 | 2.87 | 2.88 | 2.98 | 3.17 | 2.02 |
| f | 1.44 | 1.54 | 1.15 | 1.14 | 0.61 | 0 | 1.61 | 2.08 | 2.59 | 2.47 | 2.13 | 2.63 | 2.6 | 2.72 | 2.9 | 1.92 |
| g | 1.03 | 1.7 | 1.52 | 1.87 | 1.63 | 1.61 | 0 | 1.99 | 1.8 | 2 | 2.17 | 2.19 | 2.22 | 2.34 | 1.91 | 1.53 |
| h | 2.16 | 1.53 | 2.65 | 2.79 | 2.37 | 2.08 | 1.99 | 0 | 2.1 | 0.91 | 0.9 | 2.27 | 2.21 | 2.12 | 1.94 | 1.86 |
| i | 1.85 | 2.02 | 2.71 | 3.02 | 2.8 | 2.59 | 1.8 | 2.1 | 0 | 1.77 | 1.85 | 1.59 | 1.76 | 1.66 | 1.47 | 1.52 |
| j | 2.23 | 2.05 | 2.93 | 3.17 | 2.8 | 2.47 | 2 | 0.91 | 1.77 | 0 | 1.1 | 2.3 | 2.27 | 2.12 | 1.49 | 1.95 |
| k | 2.19 | 1.57 | 2.63 | 2.84 | 2.5 | 2.13 | 2.17 | 0.9 | 1.85 | 1.1 | 0 | 1.83 | 1.76 | 1.75 | 2.02 | 1.72 |
| l | 2.23 | 2.02 | 2.79 | 3.07 | 2.87 | 2.63 | 2.19 | 2.27 | 1.59 | 2.3 | 1.83 | 0 | 0.57 | 0.89 | 2.01 | 2.2 |
| m | 2.37 | 2.07 | 2.81 | 3.12 | 2.88 | 2.6 | 2.22 | 2.21 | 1.76 | 2.27 | 1.76 | 0.57 | 0 | 0.67 | 2.01 | 2.32 |
| n | 2.5 | 2.12 | 3.03 | 3.34 | 2.98 | 2.72 | 2.34 | 2.12 | 1.66 | 2.12 | 1.75 | 0.89 | 0.67 | 0 | 1.81 | 2.36 |
| o | 2.25 | 2.52 | 3.13 | 3.49 | 3.17 | 2.9 | 1.91 | 1.94 | 1.47 | 1.49 | 2.02 | 2.01 | 2.01 | 1.81 | 0 | 2.26 |
| p | 1.41 | 1.37 | 1.73 | 2.02 | 2.02 | 1.92 | 1.53 | 1.86 | 1.52 | 1.95 | 1.72 | 2.2 | 2.32 | 2.36 | 2.26 | 0 |

Root mean square distance (RMSD) between protein blocks, in angstroms. Protein blocks represent cluster centres and do not differ between RMSD- and RMSDA- based clusters.

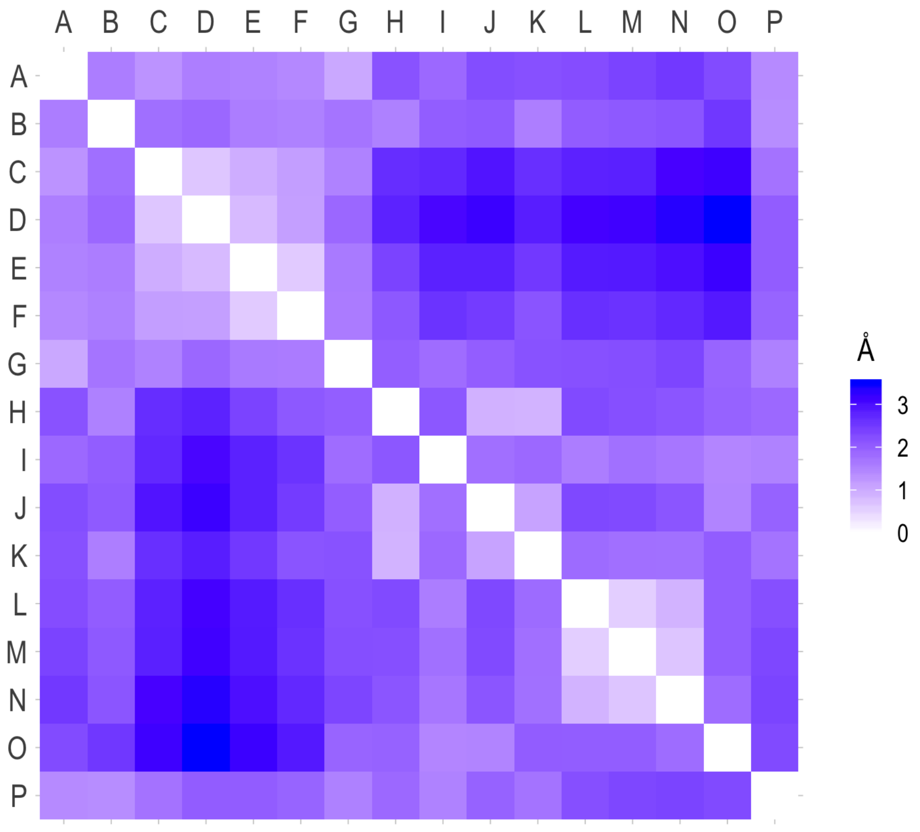


Figure S5: Heatmap of the distances between RMSD-based cluster centres.

Table S7: **Prediction accuracy: Benchmarking**

| **PB** | **Our Model** | | | | **PB-kPred** | | | **LOCUSTRA** |
|---|---|---|---|---|---|---|---|---|
| | PB frequency | accuracy | specificity | MCC | accuracy | specificity | MCC | accuracy |
| A | 5.3 | 60.8 | 97.9 | 0.599 | **67.20** | 98.15 | **0.69** | 58.16 |
| B | 5.7 | 54.4 | 98.1 | 0.569 | 52.15 | 97.72 | 0.56 | 26.14 |
| C | 4.5 | 43.1 | 97.8 | 0.439 | **58.53** | 95.95 | **0.58** | 44.81 |
| D | 12.8 | 79.9 | 94.6 | 0.712 | 67.00 | 94.12 | 0.63 | 71.58 |
| E | 3.8 | 42.5 | 98.3 | 0.448 | 57.45 | 98.96 | **0.62** | 44.74 |
| F | 5.8 | 55.0 | 98.5 | 0.601 | 60.30 | 97.21 | 0.61 | 41.45 |
| G | 6.0 | 43.9 | 98.8 | 0.540 | 43.45 | 99.19 | 0.51 | 26.84 |
| H | 1.3 | 41.2 | 99.7 | 0.514 | **61.05** | 98.82 | **0.64** | 38.45 |
| I | 3.8 | 44.8 | 99.4 | 0.578 | 59.17 | 99.18 | **0.63** | 36,87 |
| J | 1.7 | 45.9 | 99.7 | 0.590 | 49.98 | 99.40 | 0.56 | 48.19 |
| K | 5.3 | 64.6 | 98.5 | 0.661 | 63.98 | 97.67 | 0.65 | 48.46 |
| L | 4.6 | 32.1 | 98.8 | 0.405 | 59.99 | 97.69 | **0.62** | 42.71 |
| M | 30.3 | 96.5 | 82.0 | 0.726 | 75.89 | 91.03 | 0.67 | 83.76 |
| N | 3.4 | 34.2 | 99.4 | 0.474 | **62.15** | 99.05 | **0.65** | 52.08 |
| O | 2.2 | 64.5 | 99.5 | 0.695 | 63.19 | 98.70 | 0.66 | 55.1 |
| P | 3.4 | 58.2 | 98.9 | 0.602 | 59.24 | 98.25 | 0.62 | 40.8 |

We assessed the prediction accuracy of our method via leave-one-out cross-validation, meaning that a complete protein and all its 5-mer fragments were removed from the training step and predicted in the validation step. The prediction accuracy of PB-kPred and LOCUSTRA are those reported in the original articles. [18] PB frequency stand for the frequency of the fragments from the corresponding cluster in the sample, MCC stands for the Matthews correlation coefficient.[19]

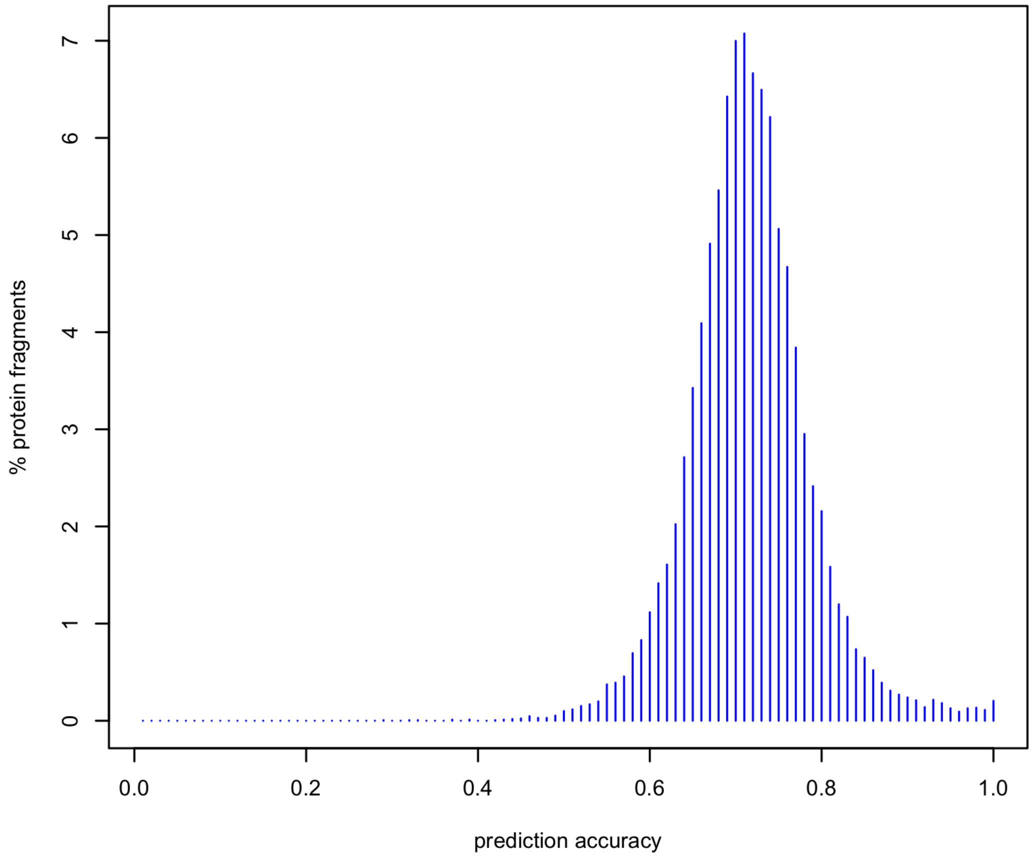


Figure S6: Distribution of the prediction accuracy calculated via cross-validation.

Table S8: **Dihedral angles of the protein blocks (de Brevern et al.)**

| PB | $\psi_1$ | $\omega_1$ | $\phi_2$ | $\psi_2$ | $\omega_2$ | $\phi_3$ | $\psi_3$ | $\omega_3$ | $\phi_4$ | $\psi_4$ | $\omega_4$ | $\phi_5$ |
|---|---|---|---|---|---|---|---|---|---|---|---|---|
| A | 41.139 | 180 | 75.529 | 13.919 | 180 | -99.799 | 131.879 | 180 | -96.269 | 122.079 | 180 | -99.679 |
| B | 108.239 | 180 | -90.119 | 119.539 | 180 | -92.209 | -18.059 | 180 | -128.929 | 147.039 | 180 | -99.899 |
| C | -11.609 | 180 | -105.659 | 94.809 | 180 | -106.089 | 133.559 | 180 | -106.929 | 135.969 | 180 | -100.629 |
| D | 141.979 | 180 | -112.789 | 132.199 | 180 | -114.789 | 140.109 | 180 | -111.049 | 139.539 | 180 | -103.159 |
| E | 133.249 | 180 | -112.369 | 137.639 | 180 | -108.129 | 132.999 | 180 | -87.299 | 120.539 | 180 | 77.399 |
| F | 116.399 | 180 | -105.529 | 129.319 | 180 | -96.679 | 140.719 | 180 | -74.189 | -26.649 | 180 | -94.509 |
| G | 0.399 | 180 | -81.829 | 4.909 | 180 | -100.589 | 85.499 | 180 | -71.649 | 130.779 | 180 | 84.979 |
| H | 119.139 | 180 | -102.579 | 130.829 | 180 | -67.909 | 121.549 | 180 | 76.249 | -2.949 | 180 | -90.879 |
| I | 130.679 | 180 | -56.919 | 119.259 | 180 | 77.849 | 10.420 | 180 | -99.429 | 141.399 | 180 | -98.009 |
| G | 114.319 | 180 | -121.469 | 118.139 | 180 | 82.879 | -150.049 | 180 | -83.809 | 23.349 | 180 | -85.819 |
| K | 117.159 | 180 | -95.409 | 140.399 | 180 | -59.349 | -29.229 | 180 | -72.389 | -25.079 | 180 | -76.159 |
| L | 139.199 | 180 | -55.959 | -32.699 | 180 | -68.509 | -26.089 | 180 | -74.439 | -22.599 | 180 | -71.739 |
| M | -39.619 | 180 | -64.729 | -39.519 | 180 | -65.539 | -38.879 | 180 | -66.889 | -37.759 | 180 | -70.189 |
| N | -35.339 | 180 | -65.029 | -38.119 | 180 | -66.339 | -29.509 | 180 | -89.099 | -2.910 | 180 | 77.899 |
| O | -45.289 | 180 | -67.439 | -27.719 | 180 | -87.269 | 5.129 | 180 | 77.489 | 30.709 | 180 | -93.229 |
| P | -27.089 | 180 | -86.139 | 0.299 | 180 | 59.849 | 21.509 | 180 | -96.299 | 132.669 | 180 | -92.909 |

Dihedral angles of protein blocks backbone used for classification and protein local structure prediction. These PB were originally designed by de Brevern et al. and used in their implementation of the PB-kPred.[9] Protein Blocks reference angles were taken from A. G. de Brevern, C. Etchebest and S. Hazout. ”Bayesian probabilistic approach for predicting backbone structures in terms of protein blocks”, Proteins, 41: 271-288 (2000)
